# Differences in running performance of single- and group-housed mice

**DOI:** 10.1101/2021.12.29.474296

**Authors:** Uma T. Plenz, Patrick O. Kanold

**Affiliations:** Department of Biomedical Engineering, Johns Hopkins University School of Medicine, Baltimore, MD, USA

**Keywords:** transgenic mouse, locomotion, behavioral phenotype, low-cost data acquisition

## Abstract

Mice are one of several common animal models in neuroscience and mouse behavior is becoming increasingly relevant. Mice are housed either in groups or alone in standard cages during which they show a variety of different behaviors. Moreover, housing conditions might alter the behavior of the mice as do social interactions. When given access to running wheels as an environmental enrichment, mice tend to run long distances preferentially during their dark cycle. However, it is currently not well understood whether and how mice utilize running wheels when single-housed or group-housed. Here we developed a low-cost running wheel data acquisition system to study running in adult C57BL/6 mice at high temporal resolution under different social conditions. As expected, adult C57BL/6 mice prefer to run in stretches during the dark cycle and mostly rest during the light cycle. When single-housed, running bouts occur independent from each other as indicated by an exponential decaying autocorrelation. In contrast, mice run ∼50% less when housed in groups of n = 3 and their temporal pattern of running exhibits a power law decay in the autocorrelation indicative of potential social interactions. Our results demonstrate that running wheels are a limited resource for which mice compete for when they are group-housed, thereby reducing their overall running activity.

**Significance Statement:** Voluntary cage wheel running is an important environmental enrichment for mice housed either alone or in groups. We show that this resource is considerably restricted for group-housed mice.

## Introduction

Mice are one of several common animal models in neuroscience and the current interest in understanding neural circuitry underlying specific behaviors makes understanding mouse behavior increasingly relevant. In experimental settings, mice are housed either in groups or alone in cages during which mice display a variety of different behaviors. Studies using environmental enrichment have shown a large influence of the housing conditions on the brain (Baroncelli et al., 2010; Kempermann, 2019; Stryker and Löwel, 2018)). Thus, housing conditions as well as social interactions might alter the behavior of mice. One of the simplest behaviors to observe and that is shared between mice and humans is locomotion.

Locomotion activity is a window into brain function as it involves motivation, control, and possibly social interaction. Locomotion activity patterns are linked to the activation of specific brain regions that are involved in the motivation to run and in controlling the amount of locomotor behavior performed (Rhodes et al., 2003). Mice are nocturnal, demonstrating a nocturnal circadian rhythm in locomotor activity. That is, they run ∼5x more during dark compared to the day (Fuochi et al., 2021; Kobayashi et al., 2020). The onset in running after dark is rapid, within 1 hr (Bains et al., 2018; Fuochi *et al*., 2021; Kopp, 2001). Observations in multiple strains of mice have shown that mice run in intermittent bouts of locomotor activity during the day (Fuochi *et al*., 2021; Kobayashi *et al*., 2020).

While bouts of locomotion are present in all mice, locomotion is enhanced when mice have access to a running wheel. In home cages without access to cage wheels, mice exhibit peak running speeds of ∼1 cm/s covering total distances of ∼50 m during the day and ∼150 – 200 m during the 12h night cycle (Iannello, 2019). The overall locomotion activity changes drastically with access to wheels. In this case, mice show a more distinct night/day activity profile compared to when they don’t have access to a wheel, while maintaining some intermittency in activity periods throughout the light/dark (LD)-cycle (Bains *et al*., 2018).

Moreover, mice run up to 100x longer distances of about 10 – 20 km/day with access to a wheel without significant increase in food consumption (Koteja et al., 1999) or major change in body weight (Zhu et al., 2021) demonstrating that running is a higly rewarding activity over alternative environmental enrichments (Sherwin, 1998) at low metabolic cost.

This physiological regulation of running in mice has genetic components. Inbred mice remain their circadian rhythm in cage wheel activity (Kopp, 2001) and running can be enhanced through breeding with females running about 3 times more distance than males (12 km vs. 4 km/day; (Koteja *et al*., 1999). Running distance varies in individual mice over days (Goh and Ladiges, 2015) and in general varies by age and strain (Goh and Ladiges, 2015). Running activity in C57BL/6 mice is similar for 4 – 12 months of age, but greatly reduced for mice > 20 months of age (Elias et al., 1975; Goh and Ladiges, 2015). Female mice tend to run longer distances than male mice, although in BL/6 strains a reverse trend has been reported with females running ∼15% less that males (Fuochi *et al*., 2021).

Running is also regulated by neuromodulators. For example dopamine release is thought to be optimized in high-running rats as both increase/or decrease of D_1_–receptor stimulation reduces running (Roberts et al., 2012). The site of dopamine action seems to be the nucleus accumbens where both the D_1_ and D_2_ –dopamine receptor stimulation selectively control wheel running and food intake (Zhu et al., 2016). Specifically, D_1_–dopamine receptor stimulation increases locomotion, whereas D_2_–dopamine receptor stimulations reduces locomotion. Dopamine release is also affected by social housing conditions (Pasquarelli et al., 2017), thus the housing conditions of mice might influence wheel running behavior.

Here we studied wheel running in mice during single and group housing. We show group housing reduces overall running activity in group-housed mice by ∼50% compared to single housing and changes their temporal organization in running. Our data suggest that running wheels provide a significant environmental enrichment of which mice compete for when housed in groups.

## Materials and Methods

All animal procedures were approved by the Johns Hopkins University and NIH Animal Care and Use Committee. We used male and female adult mice (C57BL/6J background, Jackson Labs, #000664).

We fitted round, multi-use neodymium permanent magnets (6×3 mm; e.g. Diymag) to a running wheel (Tecniplast, USA). A magnetic induction switch (Reed; contact normally open; plastic; Wowoone; 2.5mm×14mm) was positioned at the back of the wheel. Sensor data was acquired with a National Instruments data acquisition board (NI USB-6215, bus-powered M-series). Our magnetic sensor was stabilized with shrink tubing that neatly fit into the 1 ml tube of a syringe. This arrangement allowed for quick removal of the sensors during cage changes and for wheel washing/sterilization procedures between experiments. We also added an LED that conveyed the activity within each channel without the need for a computer to aid with proper positioning of the magnetic switch prior to recording. The cost for parts beyond the NI-card was <$10 per cage. Data was acquired at a 100Hz sampling rate using custom scripts in MATLAB (Mathworks).

To accustom mice to wheel running before recording, mice were group-housed and separated by sex for up to several weeks with access to a single wheel in each cage. The day of the individual locomotion assessment, animals were single-housed for up to 72 hrs with non-competitive access to a cage wheel attached to the data acquisition system. Locomotion activity was continuously recorded at a sampling rate of 100 Hz/cage wheel for up 72 hr under a 12h:12h light-dark (LD) cycle. Mice were not sound isolated from each other and were free to communicate across cages. Group running was assessed in their respective home cages.

Running activity was analyzed with custom-written code in MATLAB (Mathworks). All data is expressed as mean ± standard deviation, if not stated otherwise.

The autocorrelation was calculated from running activity during the D-phases using a standard z-normalized co-variance measure for a maximal shift of up to 500 s corrected for tapering. The reduction (R) in wheel running activity during group housing was estimated as the ratio of R=(D_Grp_ – 3·D_Sgl_)/ 3·D_Sgl_, in which D_Grp_ is the mean distance run during group housing. D_Sgl_ is the mean distance run of the corresponding 3 mice when single-housed. The ratio was expressed as percentage, i.e. multiplied by 100.

## Results

We studied voluntary wheel running of wildtype C57Bl/6 mice in two cohorts, each consisting of 3 females and 3 males housed in groups (age 3 – 5 months; weight 25 – 35 g). In order to study running with high temporal resolution we developed a low-cost system to monitor the running wheels. We instrumented a commercial cage wheel with 4 magnets and a reed switch and aquired signals at 100Hz using a data acquisition board (DAQ) board (Fig. 1). Each AD card has 16 channels, allowing us to monitor up to 16 cages. To estimate the running behavior for individual mice, mice were single housed with exclusive wheel access for 2 – 3 days followed by measurement of running activity after return to their respective home cages for several days (group housing). For data analysis, measures were averaged over several days of recordings for single-housed and group house conditions. Single-housed running activity occurred in ‘bouts’ that are abrupt onsets of running activity often at maximal speed that cease equally rapidly (Fig. 2A, B). As expected for nocturnal animals, bouts occurred mostly during the dark phase, could last many hundreds of seconds, and revealed running speeds of up to ∼120 cm/s resulting in clear peak-velocities in the speed distribution (Fig. 2C, 3A).

**Figure 1.**
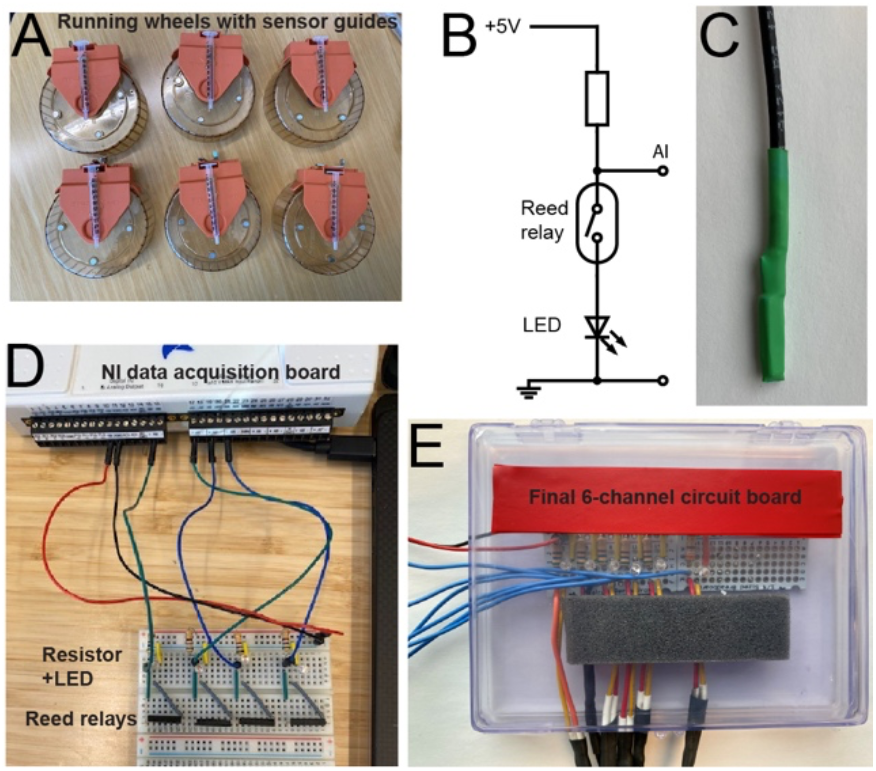
Low-cost data acquisition system to assess spontaneous wheel running (locomotion) in mice. (**A**) Plastic running wheels (Tecniplast, USA) modified for low-cost data acquisition by attaching round neodymium magnets (6×3 mm) and a 1 ml outer syringe tubing to hold the magnetic induction switch. The syringe tube is positioned such that the collar prevents mice from accessing the sensor cable. (**B**) Circuit layout connects multiple channels with magnetic induction switches (Reed) to the National Instruments (NI) data acquisition board (USB-6215). Four channels and +5 V, DC (*red*), ground (*black*) for power support are shown. (**C**) Channel circuit diagram. (**D**) Compact 6-channel circuit board with attached sensor cables (expandable to 16 channels).

**Figure 2.**
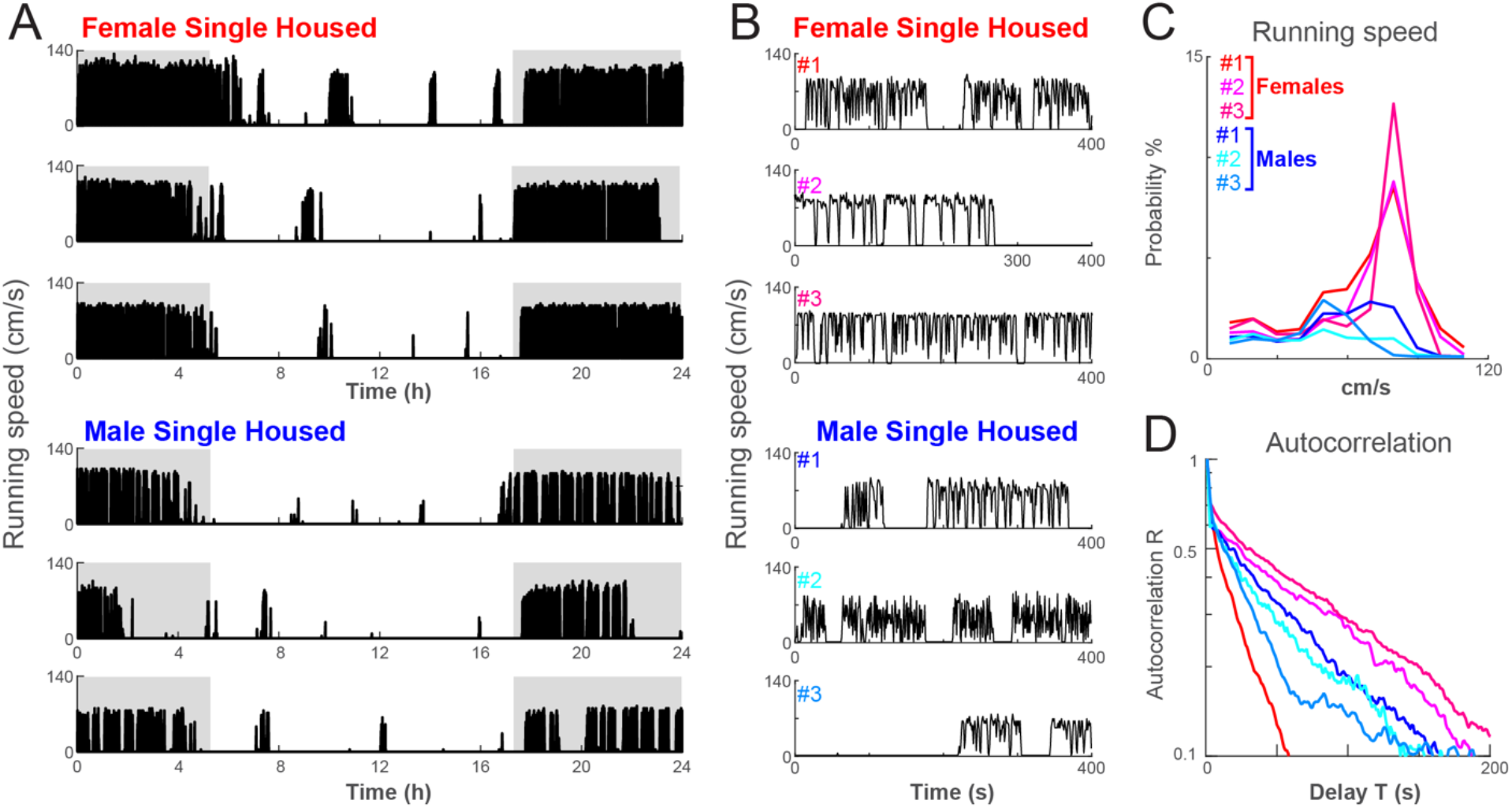
Wheel running activity in C57Bl/6 mice exhibits individual running patterns and sex differences. (**A**) Mice run predominantly during the D-phase at speeds that can reach up to 120 cm/s. During the L-phase, intermittent running bouts are observed which are not synchronized between mice (Cohort 2). *Grey*: D-phase. Simultaneous recording of n = 6 mice housed individually for one 24 h LD-cycle. (**B**) Wheel running of individual mice at higher temporal resolution. Single mice run in ‘bouts’, periods of rapid running that terminate abruptly. Note that some mice maintain consistently high running speed during bouts (row 2, 3), whereas other mice show large fluctuations in speed during bouts (row 1, 5). (**C**) Corresponding speed distributions over a full LD-cycle for single mice (for color code see *B*). (**D**) The exponential decay of the autocorrelation in running activity (semi logarithmic plot) indicates independence in the temporal occurrence of running bouts for single-housed mice (for color code see *B*).

**Figure 3.**
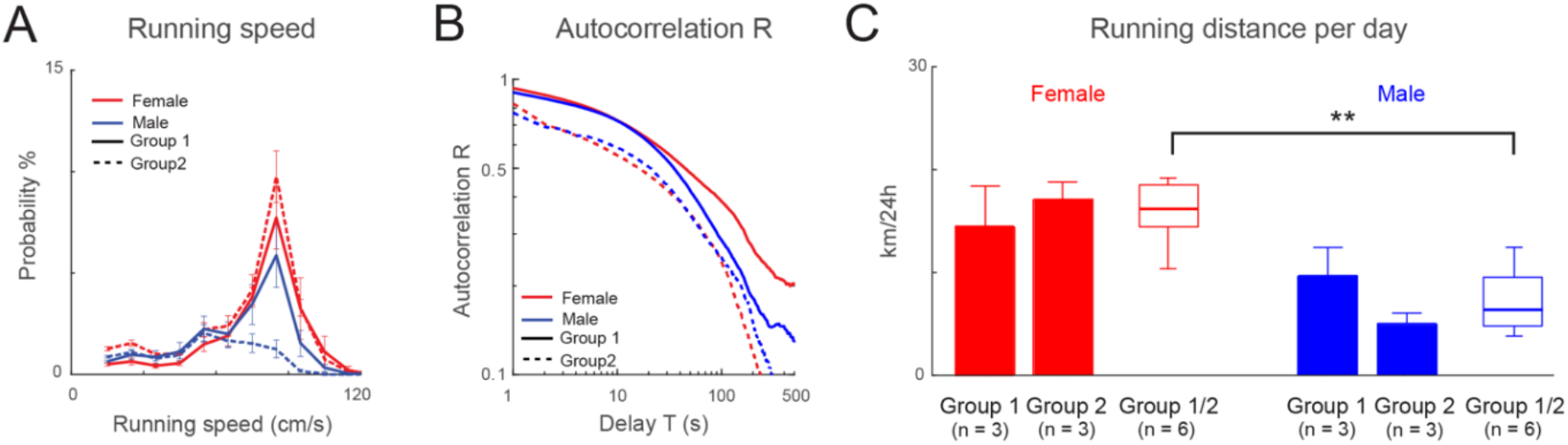
Female C57Bl/6 mice run further than male C57Bl/6 mice. (**A**) Summary in speed distributions for mice during single housing separated by cohort and sex. Females displayed similar distributions for each cohort. Running activity of male mice in cohort 1 was similar to that of female mice, whereas male mice of cohort 2 ran less. (**B**) Corresponding average decay in time course for single-housed mice separated by cohort and sex. In loglog coordinates, the exponential decay appears as a downward curved function. (**C**) Summary statistics for total distance run in single-housed mice (number in brackets indicate number of animals recorded). Note that females tend to run longer distances per day in both cohorts. Number in brackets indicate animals tested. All data are mean ± SD.

We next assessed the temporal organization of bouts by calculating the autocorrelation for the time course in running activity. If bouts exhibit a specific duration and recur at regular intervals, the autocorrelation function would be oscillatory. Alternatively, if bouts occur randomly with random durations, the autocorrelation decays like an exponential function over time. We found the autocorrelation to decay exponentially, suggesting that bouts during single housing are random occurrence of wheel running (Fig. 2D, 3B).

We next measured the distance mice covered while running. We found that female mice in both cohorts ran more than male mice reaching up to ∼10 km per 24 hr. Male mice in cohort 2 displayed the lowest running activity of ∼ 3 km per day. On average female C57Bl/6 mice covered larger distances than male C57Bl/6 mice (Fig. 3C, *p* = 0.0065; Kruskal-Wallis; n = 12), even though variability can be high.

We next investigated the effect of group housing on running patterns by housing three same sex mice together in each cage. We observed that group-housed mice also ran predominantly in bouts during the dark cycle (Fig. 4A, B). However, the temporal running patterns were different in the group housing scenario when compared to the single housing scenario. As a group, wheel locomotion increased in both female and male cages with a temporal pattern that became more continuous, though bouts were often clearly visible (Fig. 4A, B). Overall, group-housed mice maintained peak velocities of ∼80 cm/s, similar to single-housed mice (Fig. 4C-F). The increased continuous running activity suggested a change in the long-term (>100 s) temporal organization of how mice utilize the running wheel compared to when they are placed in single housing conditions. Indeed, the autocorrelation in running activity changed from an exponential distribution typical for single-housed runs (*cf*. Figs. 2D, 3B) towards a power law (Fig. 4E, F). This power law suggests current running activity to have long-lasting influence on running activity up to 500 s later.

**Figure 4.**
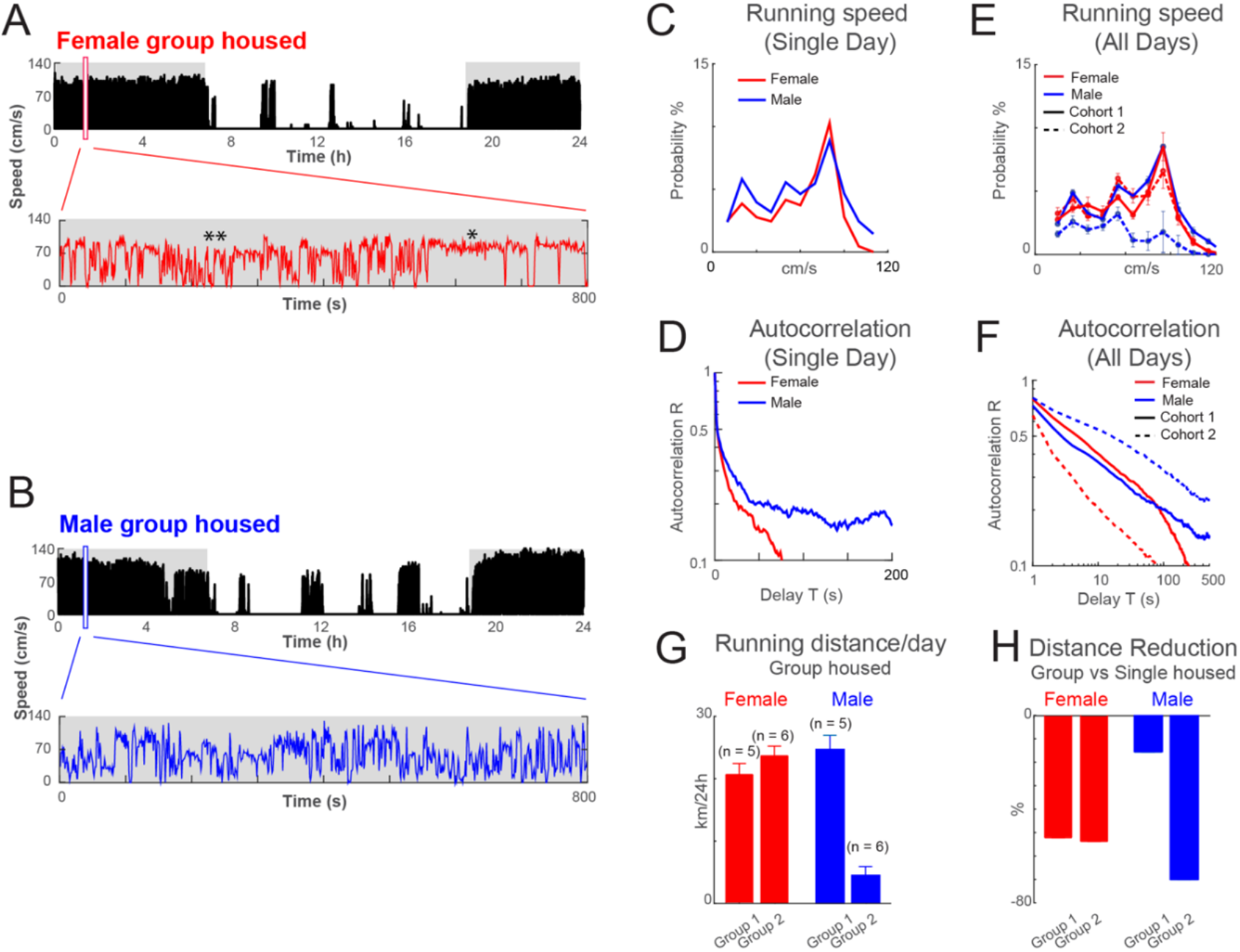
Group-housing reduces total distance in voluntary wheel running when compared to single-housing. (**A**) Group-housed C57Bl/6 mice (n = 3 mice per cage) run predominantly during the D-phase. During the L-phase, intermittent running bouts are observed which are not synchronized between cages. *Grey*: D-phase. Simultaneous recording of mice housed in groups of n = 3 separated by sex for a 24 h LD-cycle. (**B**) Enlarged temporal time course from *A* identifies segments of highly-variable running speed in male and female cages. Highly variable, low-speed epochs (**) can alternate with stable, high-speed epochs (*) as found for single-housed mice. (**C**). Similar speed distribution in group-housed male and female mice (Cohort 1; single day). (**D**) Summary in speed distribution for the two cohorts separated by group-housed males and females (n = 5 days for cohort 1; n = 6 days for cohort 2). (**E**) Slow decay in the autocorrelation for female and male cages reflects the continuous, long-term interactions between animals (semi-log plot; cohort 1; single day). (**F**) Power law decay in the autocorrelation of running activity visible as linear relationship in loglog plot. This power law relationship uncovers long-term temporal correlations during group housing (all cohorts; *cf*. exponential decay for single-housing in Fig. 2D, 3B). (**G**) Summary statistics for total distance run per day during group housing (number in brackets indicate days recorded). (**H**) Reduction (R) in running distance in group housing. All mice experience a reduction in total distance run per day during group housing of up to 70%.

We next compared the average distance run by mice when single-housed (Fig. 3C) to the total running activity of the same mice when group-housed (Fig. 4G). We found that group-housed mice on average ran between 20 – 70% less compared to when they were single-housed (Fig. 4H). We found this reduction in all groups, particularly prominent among female mice, yet also in male mice that ran the least.

## Discussion

The present study introduces a low-cost, scalable acquisition system to study voluntary wheel running of mice in their respective cages. We analyzed the temporal organization of the voluntary running of mice when having exclusive access to a wheel (single-housed) compared to when mice shared a wheel with up to 2 others of the same sex (group-housed). We found that mice on average ran up to 70% less when group-housed and their overall running patterns change from isolated running bouts at maximum speed to more continuous running activity suggesting potential social interactions. These findings suggest that access to a running wheel provides an environmental enrichment for which mice compete for.

### Comparison with alternative available cage wheel recording systems

We used a standard NI data acquisition board and added simple external magnetic switch circuits. The design was sturdy, yet flexible enough to rapidly record mice in their home cage over extended periods of time. Detailed protocols have been published for studying voluntary wheel running in mice (Goh and Ladiges, 2015) and numerous locomotion monitoring systems have been developed for mice based on various techniques such as horizontal, rotating platforms (Zhu *et al*., 2021), and video tracking (Kobayashi *et al*., 2020) including tagging individual mice with RFID tags (for review see (Bains *et al*., 2018)). Our low-cost monitoring system, in contrast to other commercially available systems that use their own caging system, can be easily implemented and flexibly employed in many animal facilities and laboratory environments. Our system uses magnetic switches which, in comparison to optical detectors, are resistant to dust. Moreover, since we utilized four magnets per wheel, we could track wheel motion with high precision allowing the detailed analysis of running bouts.

### Single- vs. group-housed running

The present study shows that single-housed mice easily run >10 km/day, yet some male mice run significantly less indicating a high level of variability even in inbred strains. Therefore, studies of running behavior benefit from low-cost monitoring systems. The finding that group-housed mice barely run a longer distance together per day suggests a significant limitation in wheel access during group housing for individual mice. The consequences of these changes and limitations during social housing are not well understood yet. Specifically, it is not clear how mice divide this limited resource during group housing. The reduction in overall running during group housing was also found in male mice that ran the least amount (Cohort 2) suggesting that this reduction does not solely arise from limited occupancy capacity of the wheel when mice are group-housed.

We found an exponential decay for the temporal organization of running bouts during single housing that changed to a power law decay during group housing. Such a profound change in autocorrelation suggests potential long-term, social interactions among mice when sharing a single wheel. The distribution in the duration of locomotion and resting times have been successfully described by exponential and power law functions respectively in both human and rodents (Anteneodo and Chialvo, 2009; Nakamura et al., 2007). Those functional differences suggest different mechanisms underlying running and resting. Indeed, changes in the power law slope have been linked to mental disease in humans (Chapman et al., 2017; Nakamura *et al*., 2007). Identifying periods of minimal locomotion requires a thorough exploration of their threshold dependency. In addition, the animal producing the current running bout needs to be identified during group housing. These technical challenges are beyond the scope of the current work. Instead, we used the robust measure of the autocorrelation, which does not require the introduction of a threshold and is sensitive to both changes in running duration and quiet periods between running bouts. Our demonstration of a significant change in autocorrelation between single and group housing supports further exploration of the underlying temporal organization in running activity. For example, simultaneous video recordings along with remote readable ID-tagging could be used to study the social hierarchy established during wheel running under those conditions. If socially stratified, running wheel occupancy under social housing conditions could be used as a proxy to identify social hierarchy among colonies. This method might be expandable to transgenic strains, which demonstrate distinct locomotion activity patterns in response to their LD-cycle (Fuochi *et al*., 2021; Kopp, 2001). Differences in locomotion activity patterns and/or wheel occupancy could be used to identify differences in social hierarchies or social communication in models of disease, changes in social hierarchies in response to specific drug treatments (Kobayashi *et al*., 2020).

## Acknowledgements

POK designed and supervised the study. UP contributed to system design, system construction, and study. UP and POK analyzed data. UP and POK wrote manuscript. We thank Dr. Peter Jendrichovsky and Amber Tietgens for help during data acquisition. Supported by NIH U19NS107474.

